# Extracting T Cell Function and Differentiation Characteristics from the Biomedical Literature

**DOI:** 10.1101/643767

**Authors:** Eric Czech, Jeff Hammerbacher

## Abstract

The role of many cytokines and transcription factors in the function and development of human T cells has been the subject of extensive research, however much of this work only demonstrates experimental findings for a relatively small portion of the molecular signaling network that enables the plasticity inherent to these cells. We apply recent advancements in methods for weak supervision and transfer learning for natural language models to aid in extracting these individual findings as 283k cell type, cytokine, and transcription factor relations from 64k relevant documents (53k full-text PMC articles and 11k PubMed abstracts). All data, results and source code available at https://github.com/hammerlab/t-cell-relation-extraction.

## Introduction

Many promising cancer immunotherapy treatment protocols rely on efficient and increasingly sophisticated methods for manipulating human immune cells. T cells are a frequent target of the laboratory and clinical research driving the development of such protocols as they are most often the effector of the cytotoxic activity that makes these treatments so potent. However, the cytokine signaling network that drives the differentiation and function of such cells is complex and difficult to replicate on a large scale in model biological systems. Abridged versions of these networks have been established over decades of research but it remains challenging to define their global structure as the classification of T cell subtypes operating in these networks, the mechanics of their formation, and the purpose of the signaling molecules they excrete are all controversial, with a slowly expanding understanding emerging in literature over time.

We present an automated method for extracting links in these networks and then apply this method to a large corpus of PMC articles relating to immunology. All of the links extracted relate exclusively to T cell subtypes and include cytokines that either induce differentiation (hereinafter referred to as “inducing cytokines”) of these subtypes or are secreted by them (“secreted cytokines”), as well as the transcription factors involved in the path along any developmental trajectory (“inducing transcription factors”). This former type of link, between T cell subtypes and secreted or inducing cytokines, was quantified over a large corpus and shared first in the immuneXpresso [1] (iX) database, which also includes a much larger breadth of cell types and covered the entirety of PubMed abstracts available at the time. A primary goal in this study is to be able to recapitulate much of this data while utilizing more expressive language models trained with less manual annotation effort, a capability conferred by recent advancements in weak supervision. We also extend these efforts to incorporate PMC full text articles, which provide access to approximately 6 times as many relations per relevant document than PubMed abstracts (5.1 relations per full-text article vs .8 relations per abstract only article when pooling across all 3 relation types), as well as transcription factors after lending credibility to the shared relation types through comparison with those first published in iX. Finally, we compare a weakly supervised approach, using Snorkel [2], for relation extraction (RE) to one based on transfer learning, using SciBERT [3], in order to evaluate what advantages, if any, arise from the auxiliary engineering effort inherent to weak supervision. In summary, this study is intended to demonstrate the viability of weak supervision for biological relation extraction in scientific literature as well as share a large database of T cell-specific cytokine and transcription factor relationships.

## Results

### Relation Extraction Validation

We trained and extracted 3 different relations with examples of each shown in **Figure 1**. A single LSTM classifier was trained for each relation and applied to a corpus as a means of producing a large number of *(cell type, cytokine)* or *(cell type, transcription factor)* tuples. Classifier predictions for two candidates in a relation were considered positive only above the threshold of 80% as it was determined using a test set with manually annotated labels that this improved the precision for each relation without restricting results to too few candidates. **Supplementary Figure 10** shows precision scores for various thresholds used to make this determination and **Supplementary Figure 11** shows high probability predictions for each relation, to better clarify how ideal relation candidates appear.

**Figure 1:**
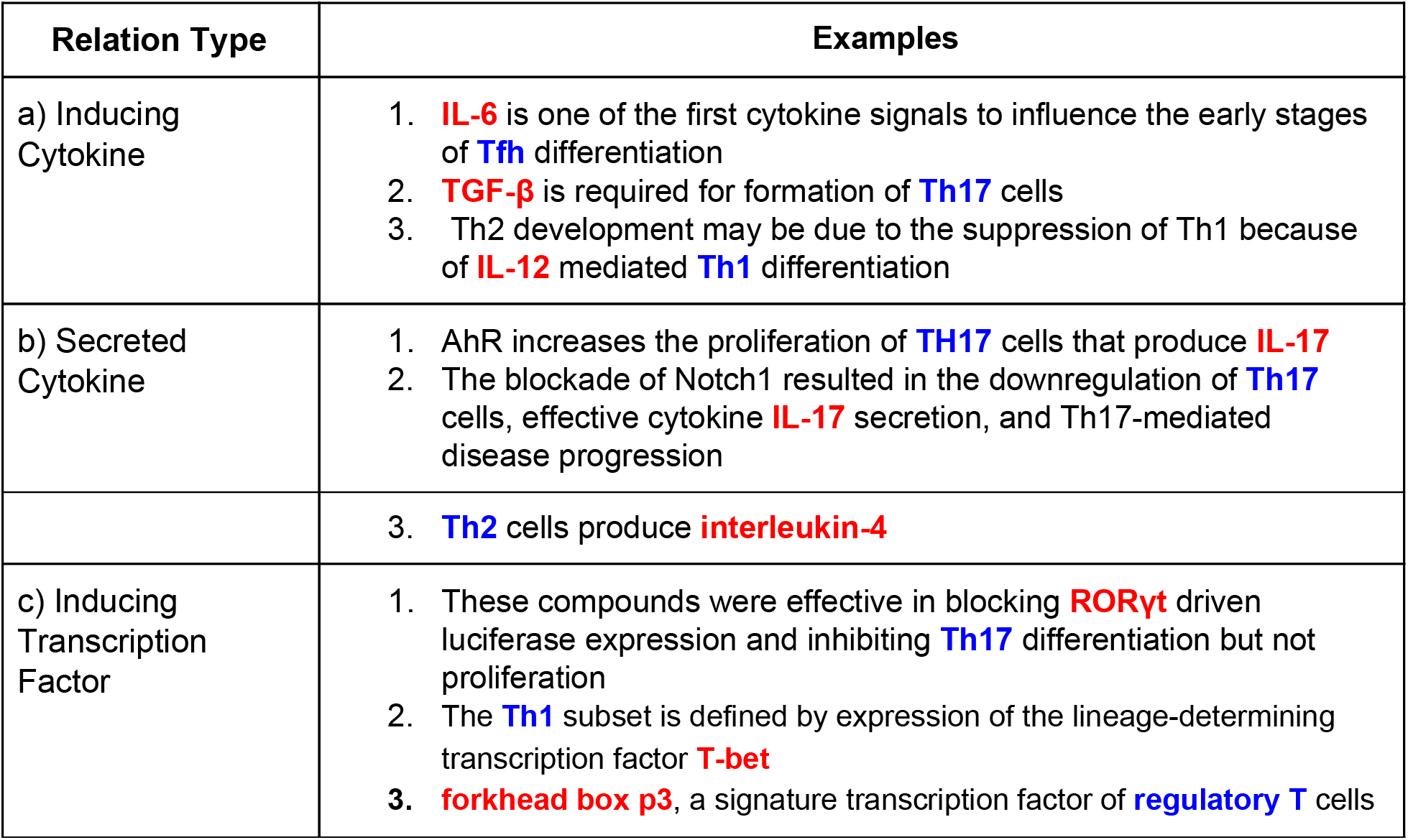
Relation example instances with an individual candidate relation highlighted as either a cytokine or transcription factor in red and a T cell subset name in blue. These examples show a few of the difficulties that arise in classifying links made through indirect language (such as in **c1**) or noun-phrases (such as “IL-12 mediated” in **a1**).

Unique document counts associated with each relation, indicating which cytokines or transcription factors are most often related to cell types, are shown in **Figure 2**. This figure demonstrates some of the breadth of information collected, which includes 75 cell types, 262 cytokines and 382 transcription factors (counts all subsequent to normalization of surface forms encountered in text). It also shows high agreement well with existing literature [4] on the cytokines most commonly associated with the function and differentiation of many T cell subtypes. See **Supplementary Figure 12** for trends in cell type mentions over time as an indication of how focus in the immunology field has shifted over the last 13 years.

**Figure 2:**
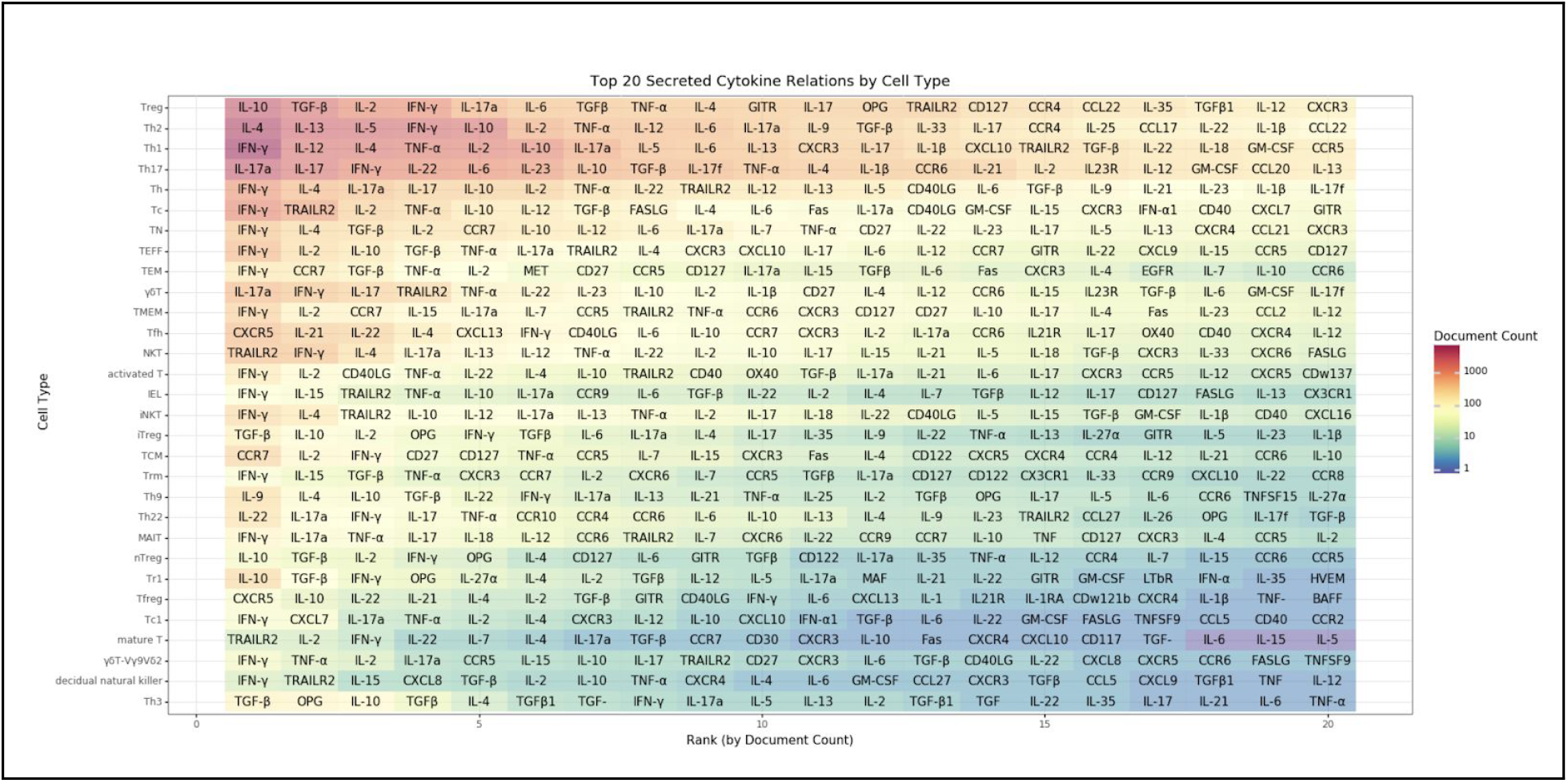
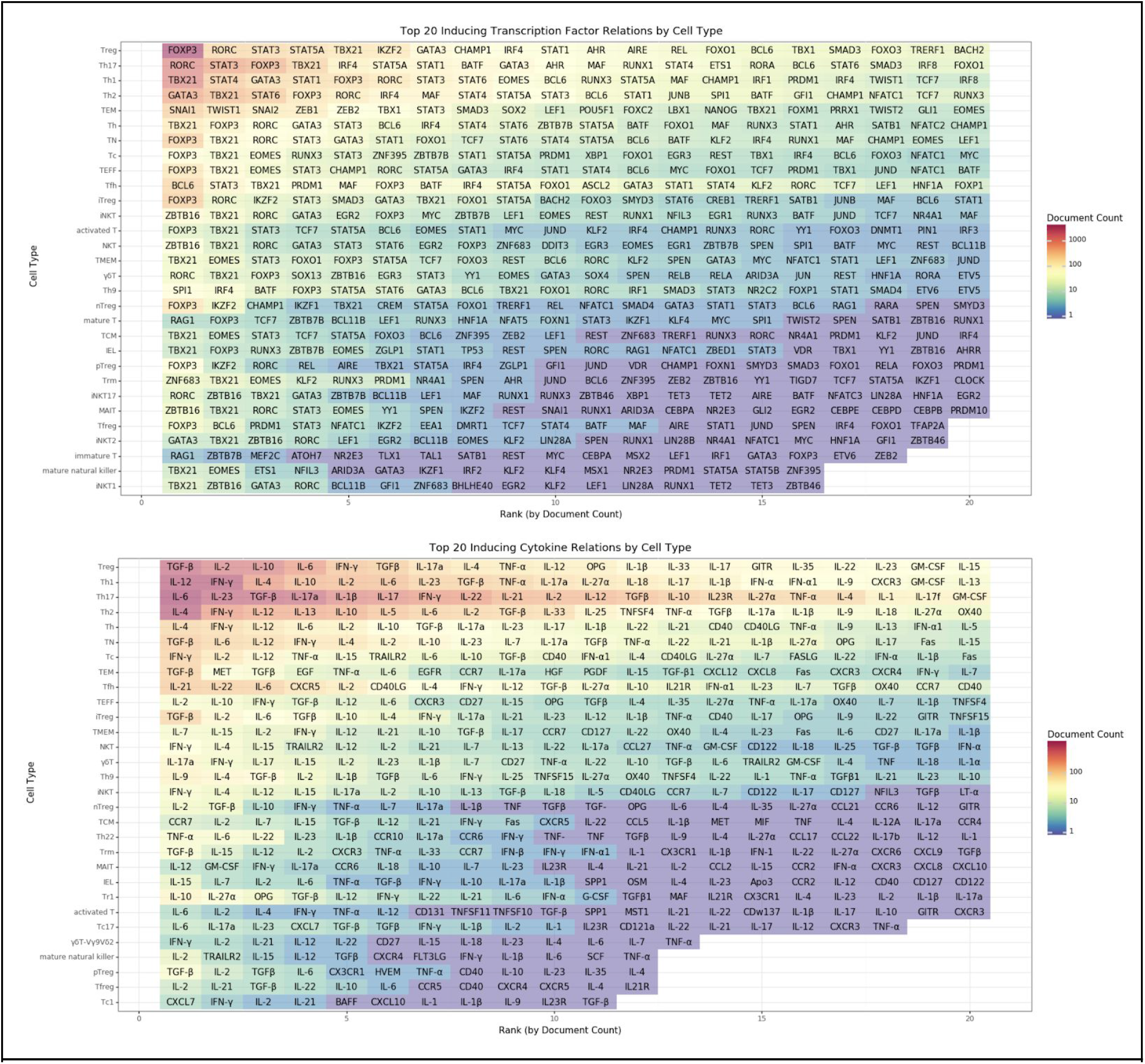
Top cytokines and transcription factors for each T cell type as determined by the number of unique documents in which a high probability (p > .8) relation instance occurs.

For further validation of T cell subtype and cytokine relations extracted, we also compared our results to those of [1], which are summarized in **Figure 3** as well as **Figure 4**. **Figure 3** shows that the number of documents associated with each relation varies based on the source (ours vs theirs) but that there is a significant level of correlation. **Figure 4** also shows alignment in the methodologies by demonstrating that conducting a similar analysis on secreted/inducing cytokine links leads to very similar results.

**Figure 3:**
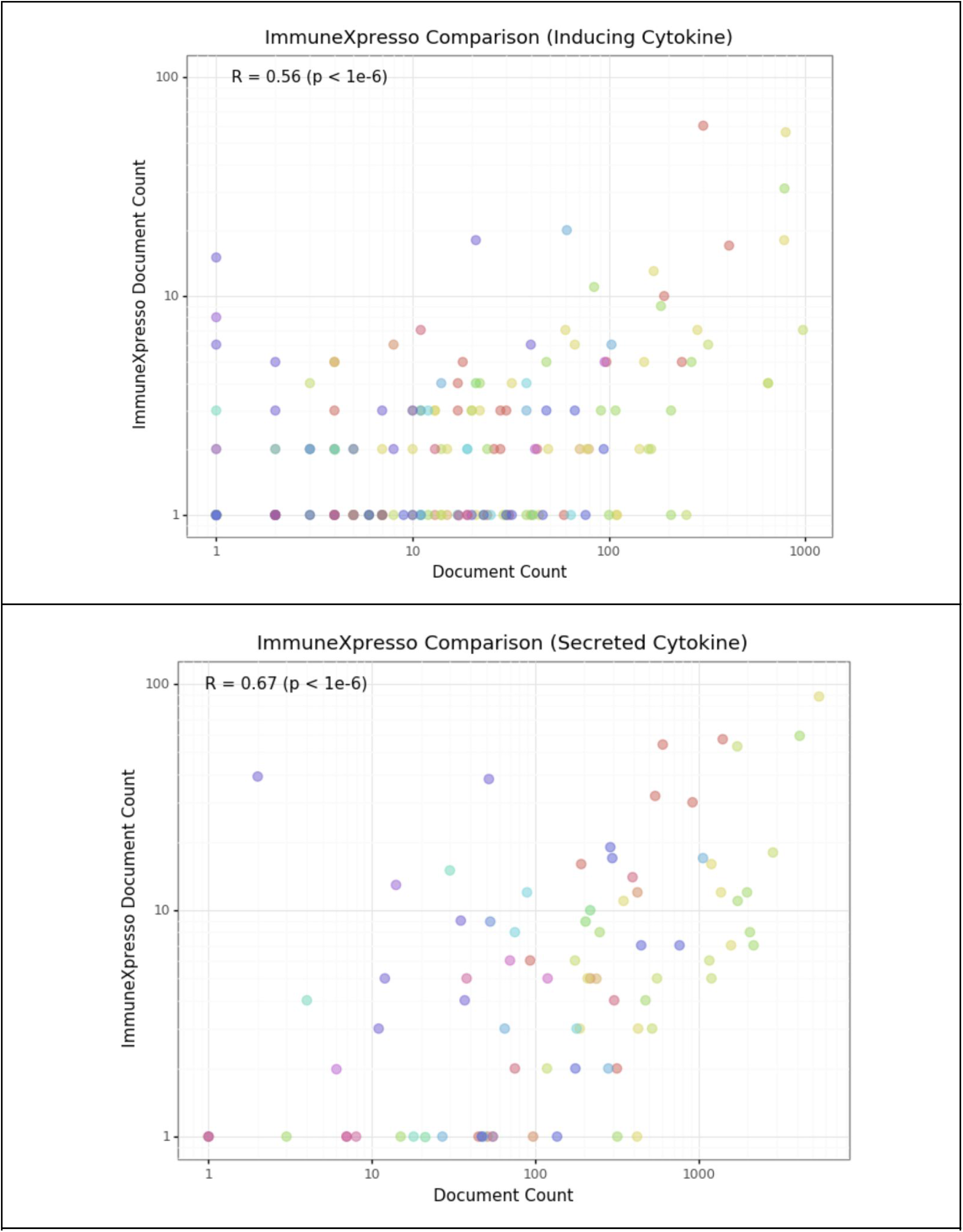
ImmuneXpresso comparison showing the number of documents discovered for each cell type and cytokine relation that was also present within this study (each dot indicates a combination of a cell type and cytokine). Pearson correlation in top left shows modest but significant agreement (two-tailed t-test).

**Figure 4:**
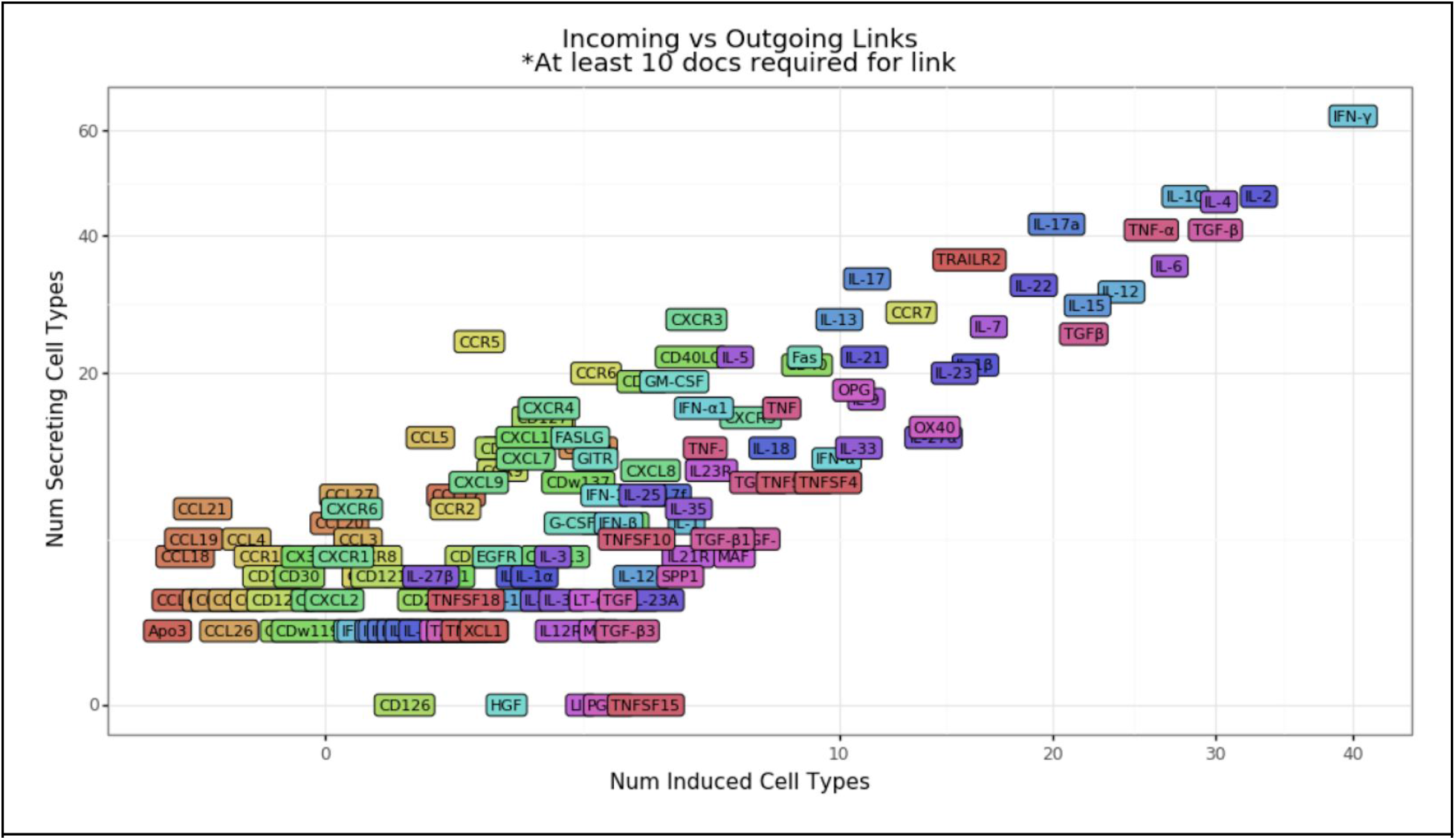
Comparison of links between cytokines and cell types that they either induce or are secreted by (or both). A single link in this case is only included when found in at least 10 distinct documents to ensure that the link is more likely to be valid. This is comparable to Figure 2c of [1] where the Pearson correlation between the number of links was found to be .86, very similar to the value of .93 found in this study.

The final, and most direct, validation of the relation extraction method included in this study is summarized in **Figure 5** where the accuracy, precision, and F1 scores for each RE task are shown for a test dataset with gold labels. This test dataset, along with a development (for evaluation of labeling functions) and evaluation dataset (for hyperparameter tuning) were all manually annotated according to these denovo <u>guidelines</u>. Candidate counts present within each manually annotated dataset can be found in **Supplementary Figure 9**. An intent of this study was to avoid manual annotation as much as possible but we found that at least ~60 man-hours (20 hours per relation type) of labeling was still necessary to build datasets large enough to evaluate and properly train Snorkel generative models, labeling functions, and standard classifiers. All annotated data and model training code is available for review or download at https://github.com/hammerlab/t-cell-relation-extraction (see the “Resources” section of the README).

**Figure 5:**
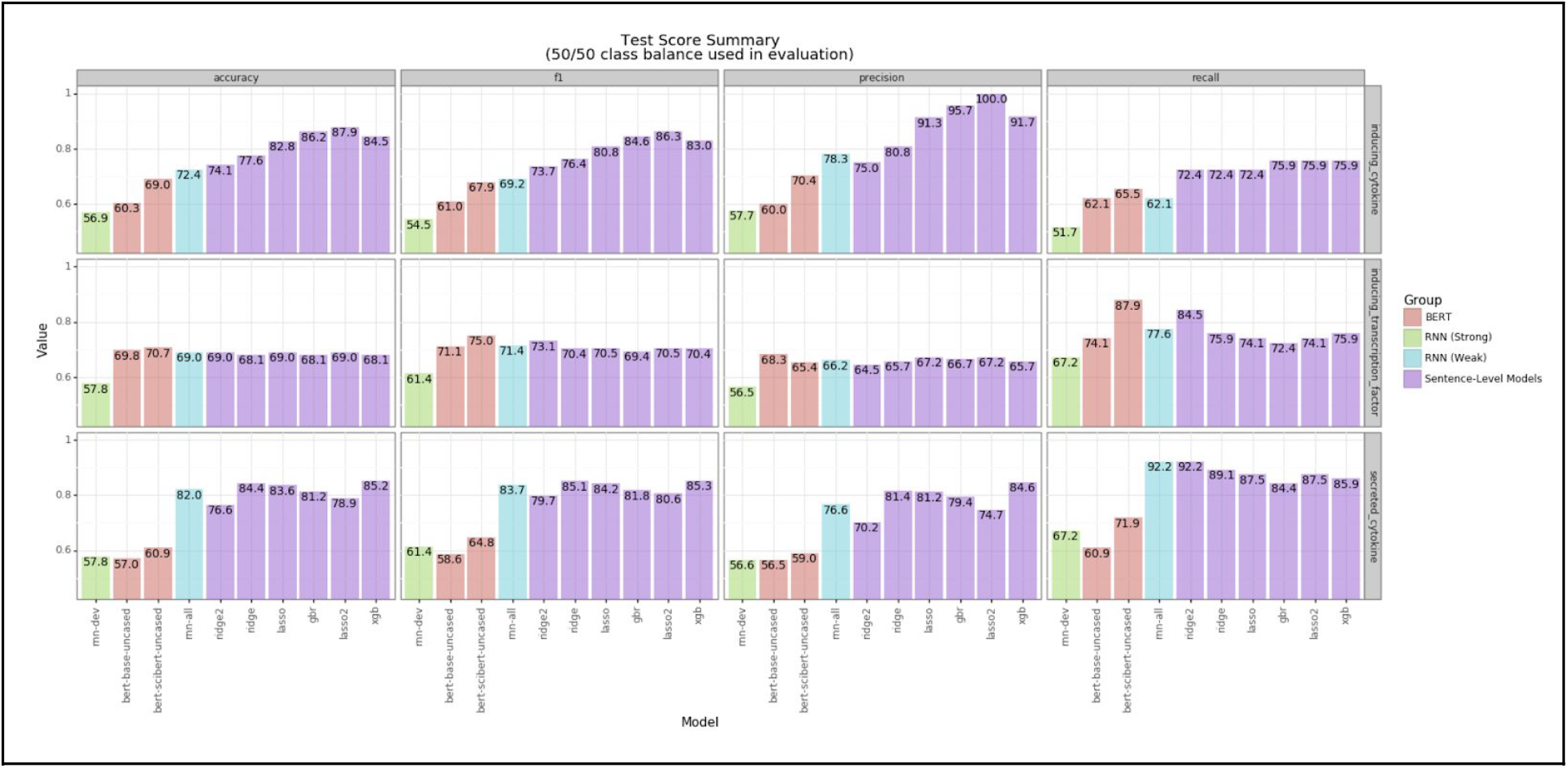
Accuracy, precision and F1 scores on test data for 4 different categories of binary relation classifiers. The “Sentence-Level Models” indicate standard machine learning models (i.e. tree and logistic regressors) applied to a set of hand-crafted features (in this case, the Snorkel labeling function outputs). The “RNN (Strong)” model indicates an RNN trained directly on the small set of examples with gold labels (~200 examples) in the development set. “RNN (Weak)” indicates the final LSTM model trained using Snorkel and “BERT” models indicate those fine-tuned on the same data as the “RNN (Strong)” model, using both SciBERT and general domain BERT for comparison.

### Expression Signature Tokenization

As a tangential finding in this study, we also explored methods for linking T cell entity mentions (in a document) to terms in an ontology but found that this is complicated by the nature of how these cells are defined and referenced. Cells such as these are often designated as belonging to particular families or types based on expression signatures, possible with a variety of assays, that capture the extent to which various intracellular transcription factors or cell surface receptor, transport channel, antigen presentation, or signaling molecules are present. An example of this would be ***“Th1 (CD4+IL-17-IFN-γhi) cells”*** which indicates both the cell type label as well as the putative markers common to that definition. Our data suggests that this somewhat redundant form is more rare than ***“Th1 cells”*** or ***“CD4+IL-17-IFN-γhi cells”*** but not so rare as to be insignificant (the ratio at which they occur is approximately 1:3:1.6 respectively). This implies that proper interpretation of these expression strings may offer a substantial improvement for entity normalization when the surface forms of the cell types are irregular. For example, ***“helper CD4+IL-17-IFN-γhi type 1 cells”*** is a synonymous reference to the example above but it is extremely unlikely to match against a large list of aliases. For this reason, we sought to determine how well these expression strings are tokenized with common tokenization tools such as ScispaCy [5]. The string ***“CD4+IL-17-IFN-γhi”*** cannot be tokenized well without knowledge of protein boundaries so we developed a method hereinafter referred to as “ptkn”, which would partition such an example as ***[CD4^+^, IL-17^-^, IFN-γ^+^]***. This method combines prior knowledge of protein aliases (from Protein Ontology [6]) as well as cytokine and transcription factor aliases with a recursive partitioning algorithm to segment character sequences with no white space into their constituent parts and normalize recognized proteins into common forms (e.g. 4-1BB and CDw137 become CD137). This approach was validated in a downstream classification task by comparing the ability of word vectors [7] resolved from this tokenization to those from the ScispaCy tokenization to capture the semantics of expression profiling (e.g. CD45RO and CD45RA are often used interchangeably with opposing signs). The results of this validation are summarized in **Figure 6** and clearly show large improvements over tokenization by regular expression, as is used in ScispaCy. **Figure 7** then shows how this tokenization can be used to correlate markers with cell types, thereby recovering common signatures. See the “Expression Signature Tokenization” section of the project README at https://github.com/hammerlab/t-cell-relation-extraction for more details.

**Figure 6:**
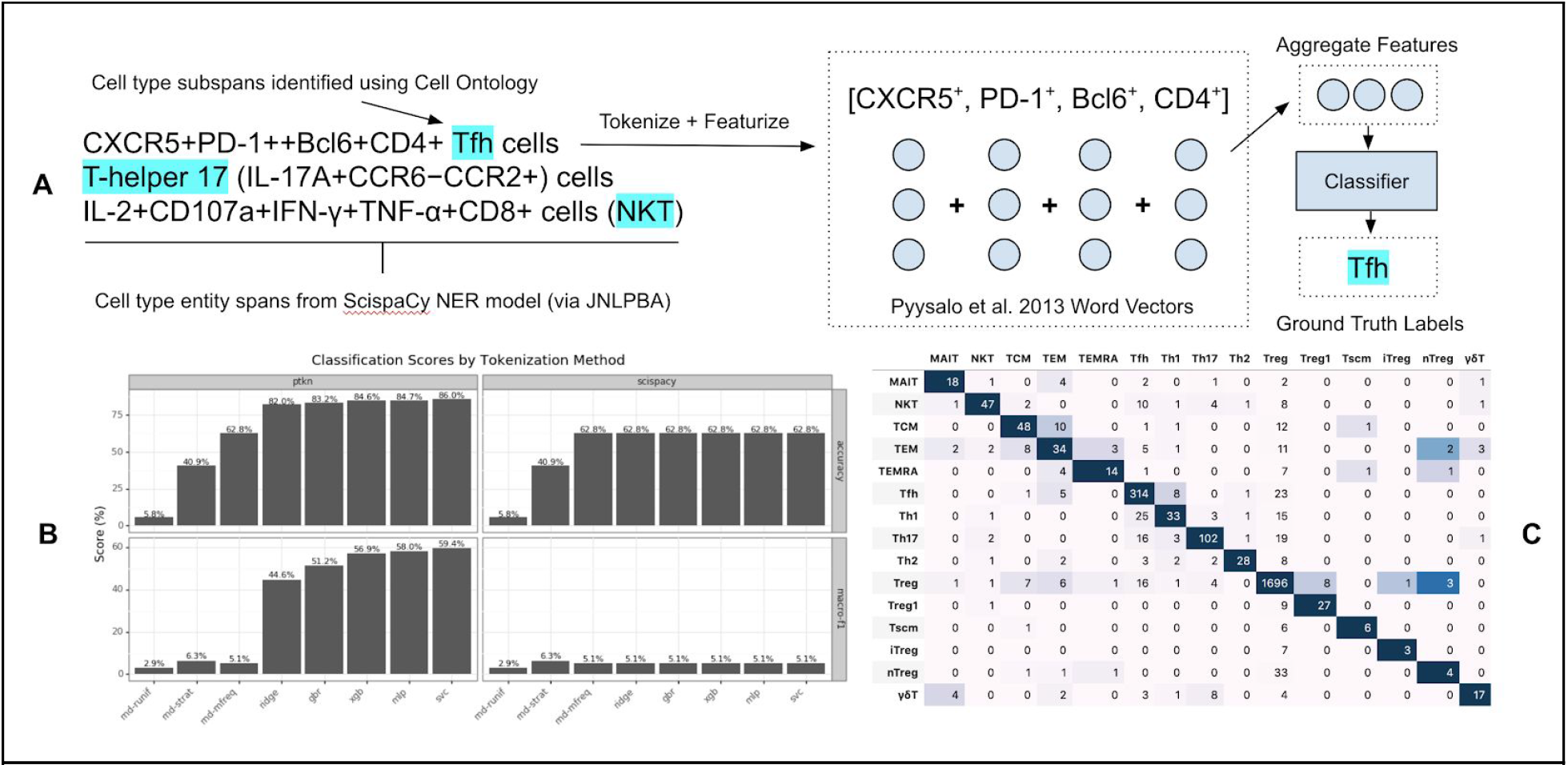
Summary of classification task used to quantify expression signature tokenization method. (A) Subspans of original entity cell type entity spans are identified that denote specific cell type labels while all other context (typically protein expression strings) is then retained to create aggregated word vectors. These word vectors are then used to classify the cell type labels with results from 10 fold cross validation shown (as averages) in (B) for both tokenization methods where all *“rnd”* estimators indicate baseline models using either random guessing *(ruinf)*, stratified guessing *(strat)* or majority class predictions *(mfreq)* and all others are standard ML estimators (support vector machines attain the highest accuracy). (C) shows the confusion matrix resulting from out-of-sample predictions in cross validation with *ptkn* tokenization.

**Figure 7:**
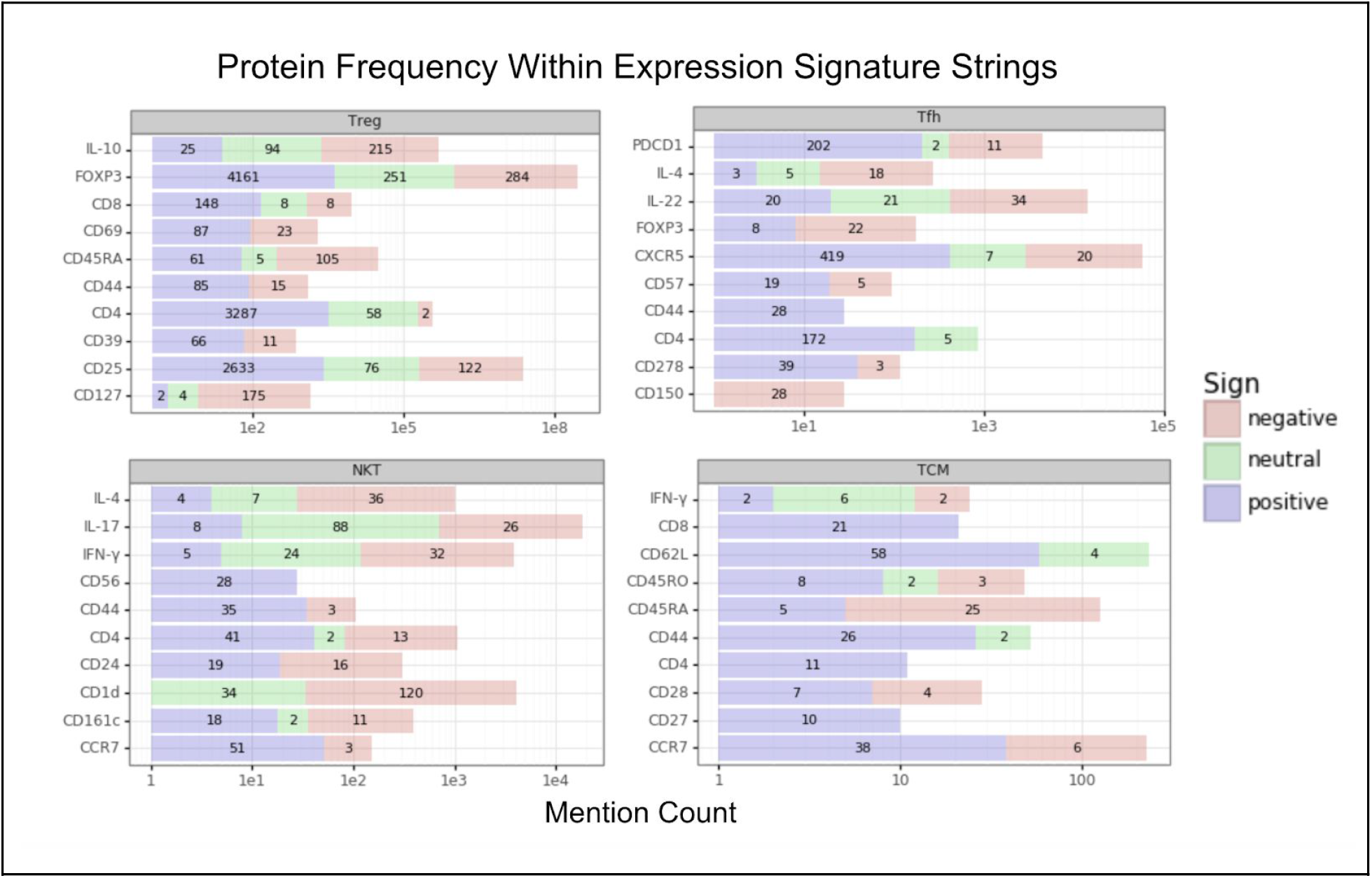
Expression signature frequency for cell spans containing signatures as well as cell type labels (e.g. *“Th1 CD4+IL-17-IFN-γ+ cells”*). The four cell types shown are Treg (regulatory T cells), Tfh (Follicular helper T cells), NKT (natural killer T cells), and TCM (central memory T cells). The “mention count” is simply the number of times that protein appeared in any expression signature string across 3,184 cell type spans containing both an expression signature and cell label, as well as at least 2 non CD4 or CD8 proteins in the signature.

## Methods

### Document Collection

Two separate corpora were collected for this study. The first (aka “Dev Corpus”) was based on a keyword query^1^ passed to the Entrez API in an effort to collect a smaller, high relevance corpus for model and entity recognition development. This query matched 120k documents, however, only 20k were included in this study with approximately 50% of those documents having full-text content available (~9.9k of 20k). MeSH terms were initially used for the queries but results sets were orders of magnitude smaller and keywords represented a higher recall method for document collection, as was appropriate for this application (for statistics on this, see this <u>Corpora Summary</u>).

The second corpus for this study (aka “Primary Corpus”) included all PMC OA articles that contained particular keywords. The keywords for this corpus were even less restrictive, limiting only to documents that contain the substrings “human”, “t cell” and “cd” (along with several variants for each). After training our extraction method on a more relevant corpus, this second corpus was intended to represent a higher recall, lower precision collection of candidate relations, ultimately containing 43.8k documents.

In both corpora, the title, abstract and body, when available, of all articles was concatenated to represent a single document for processing. 4.8k documents were present in both corpora and results from Entrez were preferred in those cases.

### Named Entity Recognition

Entity tagging for cytokines, transcription factors, and cell types was applied using known symbol and alias lookup tables^2^. Cytokine names and aliases were sourced primarily from [1] (via <u>Cytokine Registry</u>) and [8]. Transcription Factor names as well as associated aliases were integrated primarily from [8] with a large number of transcription factor aliases also provided by [9]. Cell types and associated aliases were gathered primarily from Cell Ontology [10], with all terms descending from either the T cell (<u>CL_0000084</u>) or NK cell (<u>CL_0000623</u>) terms. For all entities, a list of aliases not present in the corresponding ontologies was manually curated after identifying the need for them individually, and all aliases/names were used to identify entity spans based on exact, case-insensitive token sequence matches.

Prior to committing to this more time and labor intensive approach, the cell type NER model provided in ScispaCy (trained on JNLPBA [11]) was evaluated for this task but the recall of this model was very low on the most common of all T cell naming conventions, ThN (e.g. “Th1”, “Th3”, “Th17”, etc). An integrated approach combining the ScispaCy NER model with dictionary-based augmentations may make for a better future approach, in lieu of a NER model trained on a corpus closer to this domain.

### Training

Weak supervision for the relation classifiers was conducted as outlined in **Figure 8** where several different strategies were used to construct labeling functions. The first of these strategies included noting text patterns observed in annotation of a labeled relation set for performance evaluation. Examples of these patterns^3^ include generalized expressions of the nature of the relationship between a particular cell type and protein (e.g. “{cytokine} is (essential|important|critical) for {cell type} (development|differentiation)”). Common expressions indicative of an inhibitory relationship between a certain protein and cell type were also matched against (e.g. “{transcription factor} is critical for (suppressing|inhibiting|repressing) (induction|differentiation|development) of {cell type} cells)”). Other labeling functions included a single source of distant supervision, iX, a few structural heuristics (based on entity order or presence), dependency parse tree features intended to mirror those employed in iX and standard machine learning models (L1-regularized logistic regression, gradient boosted trees, etc.) trained on the small set of labeled dev dataset examples. Ultimately, these simpler models that utilize all other labeling function outputs as ***input*** features achieved the highest performance and proved to be an integral part of building a weakly supervised model that could surpass baseline classification performance scores.

**Figure 8:**
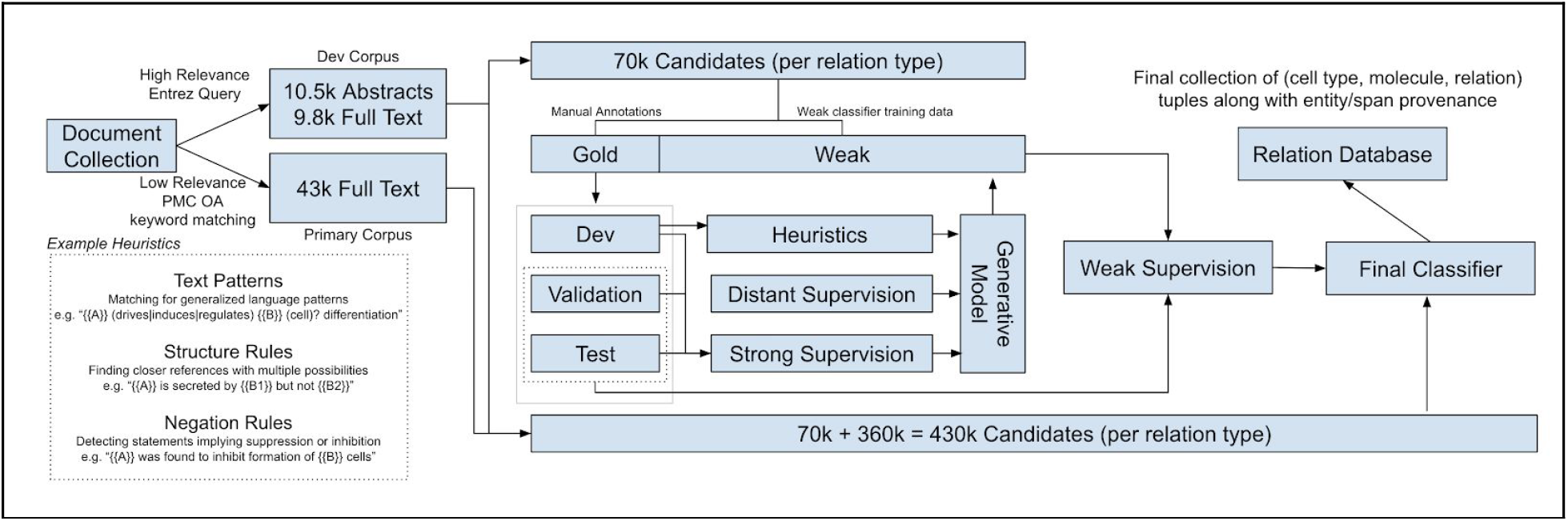
Data collection and training process summary showing approximate counts of documents and relations used to train and apply RE models.

The training dataset that was used to build the weakly supervised LSTM models for RE contained 10k examples per relation type. These datasets were then downsampled to 50% class balance based on the probabilistic labels assigned by Snorkel to smaller training sets containing ~3k examples. Hyperparameter tuning was conducted using a separate validation dataset, with gold labels, and the specifics of this grid search can be found in this <u>notebook</u>. Beyond common embedding/hidden layer sizes, dropout, learning rate, and pre-trained (using vectors from [7]) vs denovo word embedding hyperparameters, featurization strategies and the presence of positional embeddings were also included in the search. Some of the different featurization strategies included adding special tokens around the entity spans in question for a candidate as in [5] (on ChemProt RE task), using anonymized placemarks for the entities based on type (e.g. using “CYTOKINE” instead of “IL-2”), and using similar placemarks and or special enclosing tokens for entities identified in the candidate sentence but not a part of the relation in question. Positional embeddings were constructed as in [12] where each sequence presented to an LSTM is accompanied by the distance between the two entities in question for the relation as a relative offset in token count, within −127 and 127 before or after which distances are clipped to −127 and 127 respectively. This integer distance is then mapped to an index within an embedding of a configurable size (searched for in grid training) that commonly has a dimensionality between 10 and 50. For all relation types, use of position features resulted in better validation and test scores, as did dropout, decreased structural regularization (i.e. bigger models), and frozen pre-trained word embeddings. Surprisingly, featurization differences such as demarcating off-target entities and anonymizing entity spans offered little to no improvement. Hyperparameter tuning was not conducted for the Snorkel generative model itself because training with a single parameterization took ~15 minutes per relation type (w/ ~10k training examples), which made traversing any substantial fraction of the parameter space intractable. Newer versions of this implementation in Snorkel versions 0.9.x and onward may alleviate this issue, but for this study only default parameter settings were used.

## Conclusion

This work successfully reproduces a subset of functionality necessary to generate the data available in immuneXpresso (iX) while making all methods and data freely available. The relation extraction methods developed here achieve comparable precision at 83% and 92% for secreted and induced cytokine relations, respectively, while allowing for a more representative measure of recall across manually annotated relation candidates sampled without bias (i.e. candidates were not filtered using heuristics prior to annotation). This work also demonstrates that with the help of weak supervision, a single or small number of bioinformatician(s) can develop relation extraction processes with around 20 hours of annotation time expected per relation type, which may represent an improvement over the 16 authors and 11 human annotators that contributed to iX. Another advantage of this approach is that employing weak supervision as a means of training a classifier allows for the final models to be applied to text directly, rather than requiring an extensive and/or computationally expensive featurization pipeline. A disadvantage of this approach, however, is that developing accurate labeling functions was discovered difficult and time consuming for these tasks, and that even when paired with a well-tuned classifier trained on weak labels offered little improvement over far less onerous applications of transfer learning (e.g. SciBERT). Beyond our experience with applying weak supervision and transfer learning to the domain of T cell signaling network information extraction, we also share a novel set of cell type and transcription factor relations that may be valuable in the design of experiments intended to manipulate or otherwise elucidate the mechanisms of cell fate determination.

## Supplementary Material

**Supplementary Figure 9:**
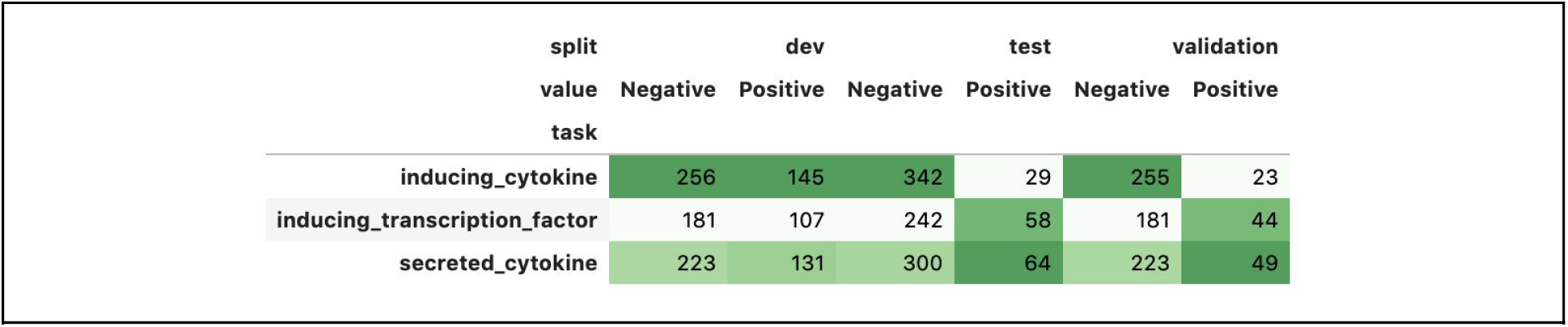
Candidate counts within each dataset with gold (i.e. manually annotated) labels. The “dev” dataset is used for developing labeling functions and building simple classification models, the “validation” dataset is used for hyperparameter tuning, and the “test” dataset is used for all final evaluations.

**Supplementary Figure 10:**
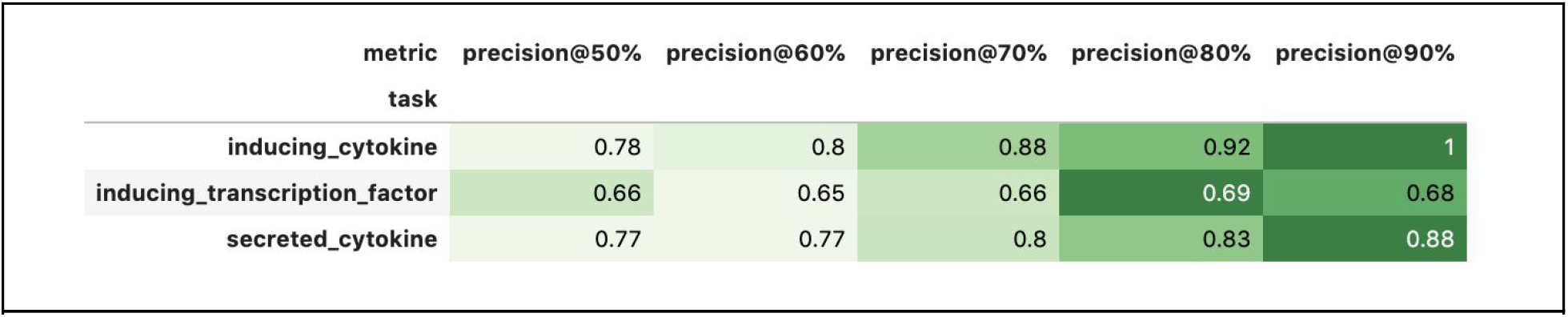
Precision scores for various thresholds on test dataset for weakly supervised LSTM model

**Supplementary Figure 11:**
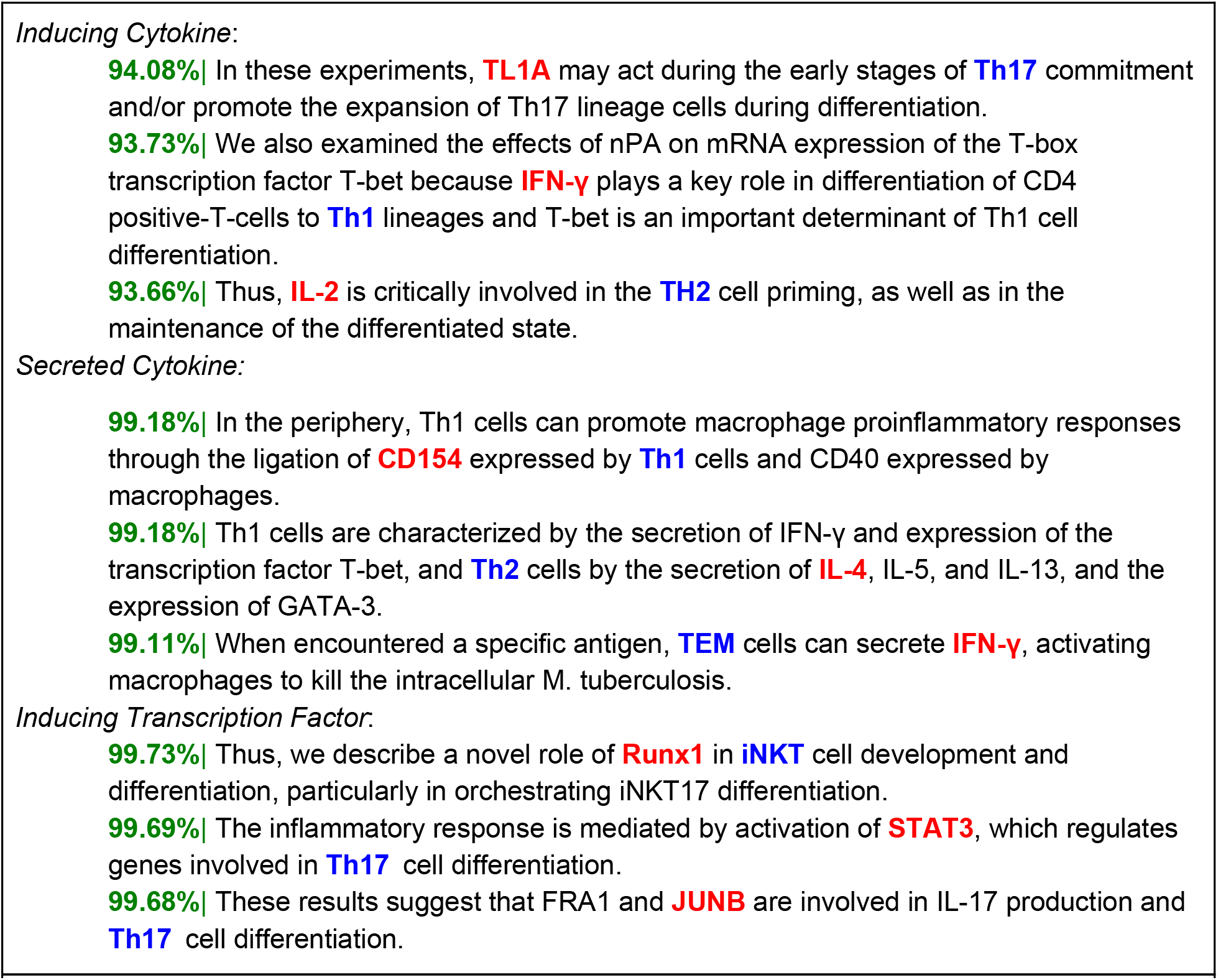
Example high probability relation classifications for each relation type.

**Supplementary Figure 12:**
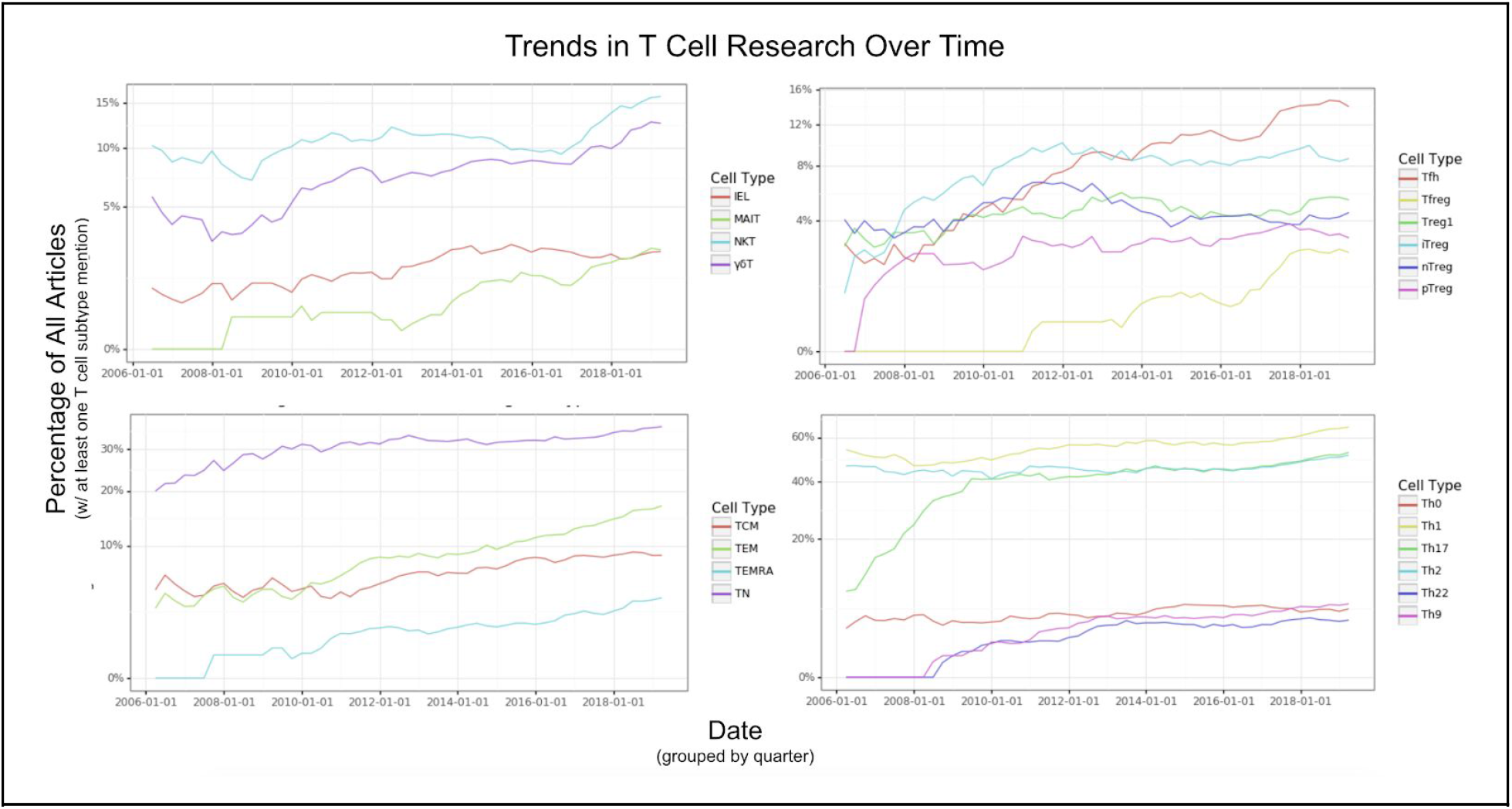
Trends in mentions of particular T cell types over time (previous 13 years) shown as percentages of 7,638 documents mentioning at least one T cell type included in this study. All values are shown as rolling averages over an 8 quarter (2 year) period. Note that regulatory T cells, and the closely related Th17 cells, have received a large increase in attention. The discovery and acceptance of now putative cell types such as follicular regulatory T, Th9, Th22, and EMRA (effector memory cells re-expressing CD45RA) T cells is also observable within this time span.

## Footnotes

1 *PMC article query: (human) AND ((t cell) OR (t lymphocyte)) AND (cytokine) AND ((differentiate) OR (differentiation) OR (differentiated)) AND ((polarization) OR (polarize) OR (induce) OR (induction)*

2 These tables are available at https://github.com/hammerlab/t-cell-relation-extraction/tree/master/data/meta

3 All patterns applied can be found at https://github.com/hammerlab/t-cell-relation-extraction/blob/master/src/tcre/supervision.py

